# The hidden land use cost of upscaling cover crops

**DOI:** 10.1101/2020.02.13.947994

**Authors:** Bryan Runck, Colin K. Khoury, Patrick M. Ewing, Michael Kantar

**Affiliations:** GEMS Agroinformatics Initiative, University of Minnesota, Twin Cities, St. Paul, Minnesota, USA; Department of Computer Science, University of Southern California, Los Angeles, CA, USA; International Center for Tropical Agriculture (CIAT), Km 17, Recta Cali-Palmira, Apartado Aéreo 6713, 763537 Cali, Colombia; Saint Louis University, Department of Biology, 1 N. Grand Blvd., St. Louis, MO, 63103, USA; Department of Agronomy and Plant Genetics, University of Minnesota, St. Paul, Minnesota 55108; Department of Tropical Plant and Soil Science, University of Hawaii at Manoa, Honolulu, HI, USA

## Abstract

Cover cropping is considered a cornerstone practice in sustainable agriculture; however, little attention has been paid to the cover crop production supply chain. In this Perspective, we estimate land use requirements to supply the United States maize production area with cover crop seed, finding that across 18 cover crops, on average 3.8% (median 2.0%) of current production area would be required, with the popular cover crops rye and hairy vetch requiring as much as 4.5% and 11.9%, respectively. The latter land requirement is comparable to the annual amount of maize grain lost to disease in the U.S. We highlight avenues for reducing these high land use costs.

## The opportunities and challenges of upscaling cover crops

Cover crops are commonly included in strategies aimed at increasing the sustainability of agricultural production systems (Figure 1). Grown between the harvest and next planting of cash crops, cover crops improve soil retention^1^, weed control^2^, soil physical properties^3^, carbon sequestration^4^, biocontrol services^5^, water quality^6^, and nutrient cycling^7,8^. They are increasingly common: from 2012 to 2017, US cover cropped area reached 6.2 million ha, a 50% increase^9^. This is due in part to large and coordinated investments by universities, nonprofits, and industry, which are improving and promoting the wider adoption of cover crops through research, advocacy, education, and outreach^10^. In spite of this uptick in adoption, just 1.7% of U.S. farmland currently incorporates a cover crop, indicating that the strategy does not yet have widespread impact in commodity crop production systems^9^. Recognizing this potential for growth, we step back from the field-scale benefits of cover cropping, and instead consider what infrastructure would be needed to plant cover crops widely across U.S. production areas, and what barriers remain to achieving this scale.

**Figure 1.**
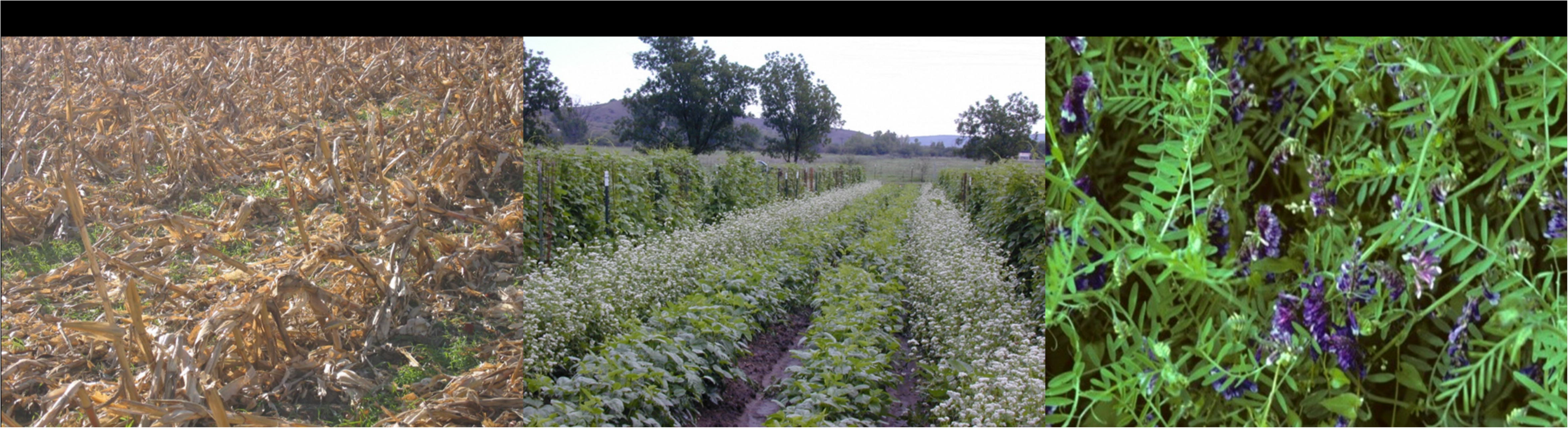
Pictures of common cover crops. **a** Cereal rye grown as a cover in corn residue in southern Minnesota (photograph by Michael Kantar), **b** Arizona bean field with cover crops of buckwheat and cowpeas intercropped between bean rows (Photo by Todd Horst), **c** Hairy vetch grown as a cover crop in southern Minnesota (photograph by David Hanson).

Perhaps the most fundamental need for upscaling cover crops is a robust seed industry that can provide an affordable, quality input for producers. Growing cover crops for seed in temperate agroecosystems usually requires foregoing production of traditional cash crops on the same land in the same year. This is because current cover crop species require most of a temperate growing season to reach reproductive maturity. As a result, widespread cover crop adoption would likely require significant arable land allocation for seed production, potentially forcing the conversion of farmed lands from cash crops, pasture, or natural systems to cover crop seed production (Figure 2). The potential scope and implications of such land use changes have not been quantified.

**Figure 2.**
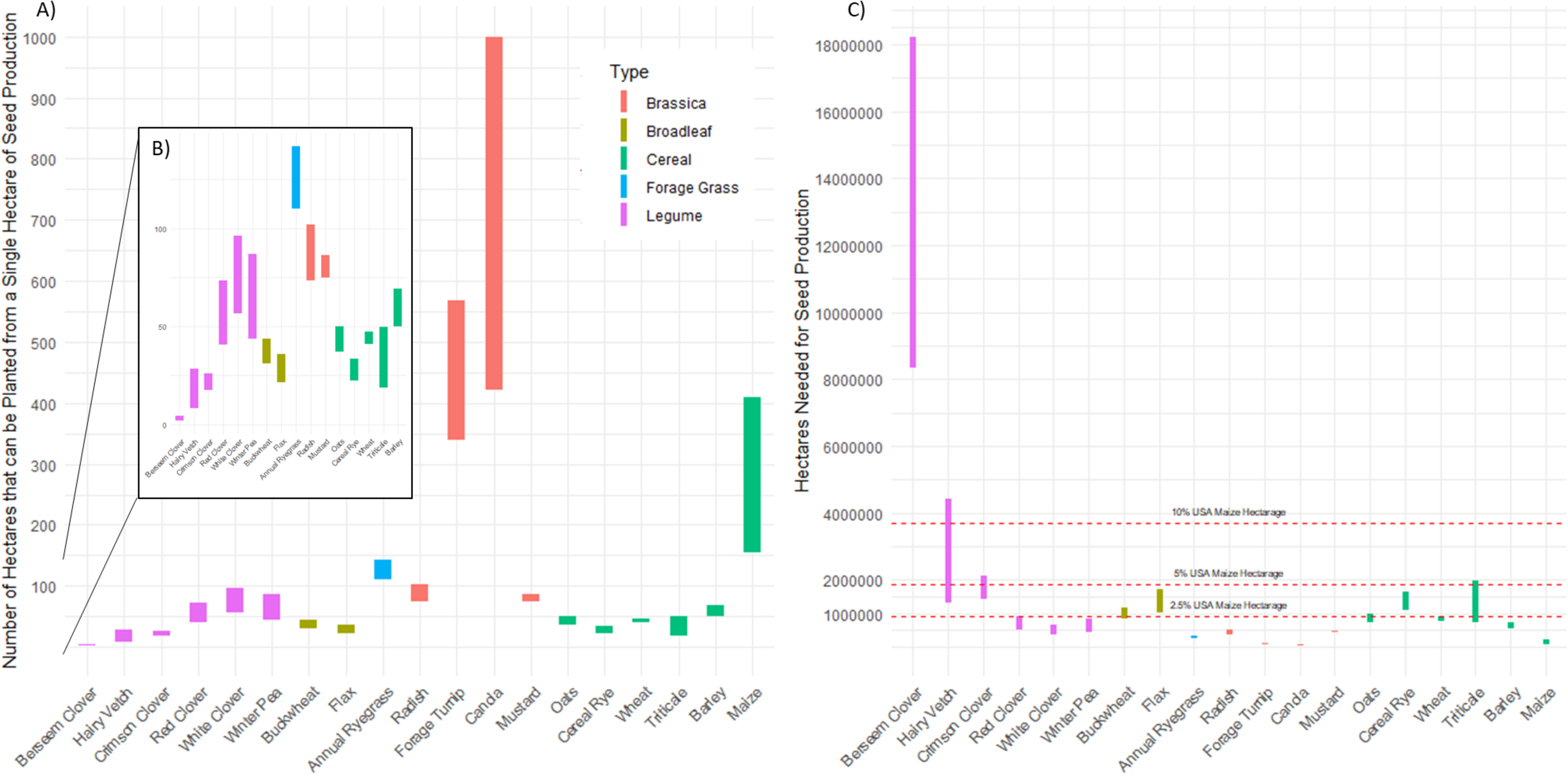
Seed production data for common cover crops. **a** Range of seed production potential from a single hectare based on commonly reported cover crop yields and seeding rates in the published literature and USDA extension **b** Zoom in of low seed yield cover crops **c** Range of area needed to support seed production based on commonly reported cover crop yields and seeding rates in the published literature and USDA extension literature. Estimates are for areal extent across the United States.

Therefore, we ask: how much land would cover crop seed production require if cover cropping was adopted widely across a major cash crop production area, such as the 37 million ha devoted to U.S. maize production? To answer this question, we compiled seed yields and seeding rates for 18 different cover crops from state yield trials, published literature, commercial seed catalogs, and farmer bulletins (Supplementary Data 1). These cover crops are marketed as suitable for use in the U.S.^11^. For each cover crop, minimum and maximum seed yield per hectare and seeding rate per hectare were used to bound the area that could be cultivated with the cover crop from a single hectare of seed production (Figure 2A), as well as the total number of hectares needed for seed production of the cover crop so as to plant the entire U.S. maize cropland (Figure 2B).

Assuming that the total maize hectarage does not change for any reason inherent to this transition, we find that the land requirements for production of cover crop seed would be on average 1.4 million hectares (median 746,000 ha), which is equivalent to 3.8% (median 2.0%) of the U.S. maize farmland. Rye (*Secale cereale* L.) – a midrange seed yielding cover crop and one of the most commonly used in the corn belt, would require as much as 1,661,000 hectares (4.5% of maize farmland), while hairy vetch (*Vicia villosa* Roth) – the lowest seed yielding – would require as much as 4,415,000 hectares (11.9% of maize farmland).

## Cover crop seed production scenarios

For the sake of illustration, we introduce two hypothetical scenarios for land use conversion for cover crop seed production, with the caveat that these scenarios do not consider all variables that go into real-world upscaling of seed production for covers. In scenario one, we consider direct competition of land between maize production and cover crop seed production and assume no change in yield due to cover cropping. If based on 2019 average maize yield data we converted land used for maize production to cover crop seed production, rye seed production would result in as much as 16,459,200 MT of maize grain removed from the market, while hairy vetch seed production would result in as much as 43,525,440 MT of grain removed. This larger number is comparable to the annual amount of maize grain lost to disease in the U.S. in 2015, which amounted to 13.5% of total production^12^.

To avoid the tradeoffs caused by producing cover crop seed on current cash crop lands, alternatives may be proposed. This caused us to consider a second scenario, where cover crop seed might instead be grown on land held in the conservation reserve program (CRP), which pays farmers to restore marginal or ecologically sensitive land to native habitat^13^. Cover cropping the entire U.S. maize area would require the equivalent of as much as 18% (rye) to 49% (hairy vetch) of the 2019 CRP enrollment for cover crop seed production^14^. Using this much CRP land to produce cover crop seed would significantly disrupt the program’s conservation and ecosystem services benefits. While further study would be needed, it seems unlikely that CRP or other marginal lands could be used instead of cash crop land to grow cover crop seed without significant ecological tradeoffs.

Acknowledging that our simplified scenarios are subject to variation in real agricultural systems, they make clear the potentially large hidden land requirements of bringing cover crops to scale. U.S. maize seed production takes less than 0.5% of the land devoted to the crop, while from our available data, the higher yielding cover crop values would still take an average of 12 times (median 7 times) as much land. This comparison is worthwhile because it makes concrete the abstract idea of cover crop seed yield by benchmarking to a well-established, efficient seed production system. Additionally, among the covers examined there was large variation (berseem clover as low seed yield; turnip and canola as high seed yield), it is important to note that ecosystem benefits of covers are not equal, and do not fit into a wide array of production systems. Hence ecologically and agronomically, it is preferable to plant rye or vetch over turnip, even though turnip has high seed yield^15^.

Planning for and mitigating projected land use needs for cover crop seed production may help pre-empt social conflicts over how to enhance agricultural sustainability^16^, which have included such high-profile disputes as food versus biofuels^17^. For example, arable lands (e.g. pasture) in other temperate regions that are not currently critical to food production could potentially be converted to cover crop seed production without major environmental cost, and in doing so may provide new market opportunities to farmers. While this could increase opportunities for participatory agronomy, it would also likely alter ecological services through changes in management intensity.

The driver behind this potential land use impact is low seed yield, acknowledging that yield estimates are highly uncertain. The United States Department of Agriculture does not keep statistics on cover crop seed yields, and agronomists researching these crops rarely report seed yields in the formal literature because the crops are most often terminated before maturity. This forced us to search for seed yield estimates in non-academic and private sources (Supplementary Data 1). Improving yield appraisals is readily achievable and would significantly improve assessments of land needed to produce cover crop seed. Yet, despite their uncertainty, these data highlight that most cover crops are almost certainly “underdeveloped” cultivated species in comparison to the generally much higher seed yields of cash crops of similar taxonomic backgrounds. Decreasing this breeding gap should reduce land use impacts of cover cropping.

Our results suggest that cover crop breeding research should shift to include more emphasis on increasing seed yield, in addition to environmental outcomes. Only a handful of cover crops are actively being bred for seed productivity (e.g., pennycress and camelina^18^). Most breeding has focused on ecosystem service values^9^ and forage quality^11^. Fortunately, advanced breeding techniques, public-private partnerships, and participatory, farmer-inclusive breeding practices could make it possible to increase the tempo of plant breeding and the subsequent adoption by farmers^19^. In particular, breeding might focus on classic domestication syndrome traits such as non-shattering, lack of dormancy, and flowering time^20^. Most of these traits have a well-known genetic basis^21,22^. Leveraging these known traits to improve seed yields may reduce land use impacts, provide economic benefits to seed producers, and improve farmers’ access to cover crop seed.

One potential way to speed the achievement of breeding goals could be to explore using a CRISPR/Cas9 approach to improve specific domestication traits, while still selecting for characteristics complementary to improved ecosystem service production. Rapid domestication using CRISPR/Cas9 recently has been successful in other plant species^23^. Specifically, the CRISPR system has been used to modify traits such as flowering, fruit size, fruit shape, plant architecture, and nutrient content in both domesticated and wild species^24,25^. However, a major limitation will be developing tissue culture protocols for cover crops as this has not been done and large variation exists in regeneration ability within and across species. In addition, potential regulation of these technologies in some world regions could translate into higher costs for producers.

## Next steps: targeting cover crop research investments

If cover crops are to be widely planted, our analysis suggests that land use for cover crop seed production could have large and poorly understood economic, environmental, and food production impacts. While the above scenarios were primarily illustrative, they highlight two research questions that require immediate attention in order to upscale cover cropping: 1) to what extent does common agronomic knowledge actually represent the yields achieved by cover crop seed growers? And 2) if seed yields for cover crops are as low as current data suggests, to what extent can we leverage breeding to increase seed yields while simultaneously improving or at the least maintaining the fertility and other ecosystem service benefits of cover crops? The answers to these questions may help indicate whether cover crops, a commonly proposed fundamental tool for sustainable crop production, will be able to upscale for widespread adoption.

## Methods

### Areal extent of seed production calculation

To identify the minimum and maximum number of crop production hectares an individual hectare of seed production could provide seed for, we divided minimum and maximum seed yield per hectare by seeding rate per hectare. We then divided the total U.S. maize hectares from the National Agricultural Statistics Service (2019) by this value to calculate the total minimum and maximum hectares needed for cover crop seed production. Full data and references for the data are available in Supplementary Data 1.

## Data Availability

All data generated or analyzed during this study are included in this published article (and its supplementary information files).

## Author Contributions

B.R, C.K.K., P.E., M.B.K. conceptualized the idea and wrote the main manuscript text and MBK prepared figures. All authors reviewed the manuscript.

